# Designed allosteric biosensors for engineered T cell therapy of cancer

**DOI:** 10.1101/2025.03.14.643388

**Authors:** Jan A. Rath, Lucas S.P. Rudden, Nazila Nouraee, Tiffany Que, Christine Von Gunten, Cynthia Perez, Flora Birch, Yashashvi Bhugowon, Andreas Fueglistaler, Aisima Chatzi Souleiman, Patrick Barth, Caroline Arber

## Abstract

Adoptive cell therapy with chimeric antigen receptor (CAR) T cells has transformed standard-of-care for selected hematologic malignancies, but relapses are frequent and efficacy against solid tumors remains limited^1,2^. The tumor microenvironment (TME) plays a key role in tumor progression^3^, and both soluble and cellular TME components can limit CAR-T cell function and persistence^4^. Targeting soluble TME factors to enhance anti-tumor responses of engineered T cells through chimeric receptors is not yet broadly explored due to the unpredictable signaling characteristics of synthetic protein receptors. Here we developed a protein design platform for the *de novo* bottom-up assembly of allosteric receptors with programmable input-output behaviors that respond to soluble TME factors with co-stimulation and cytokine signals in T cells, called T-SenSER (**T**ME-**sen**sing **s**witch receptor for **e**nhanced **r**esponse to tumors). We developed two sets of T-SenSERs targeting vascular endothelial growth factor (VEGF) or colony stimulating factor 1 (CSF1), that are both selectively enriched in a variety of tumors. Combination of CAR and T-SenSER in human T cells enhanced anti-tumor responses in models of lung cancer and multiple myeloma, in a VEGF or CSF1-dependent manner. Our study sets the stage for the accelerated development of synthetic biosensors with custom-built sensing and responses for basic and translational cell engineering applications.

---

Adoptive T cell therapies for cancer using T cells engineered with chimeric antigen receptors (CAR-T cells) are at the forefront of clinically applied synthetic immunology^1,2,5^. Direct cytotoxic targeting of tumor cells by adoptive transfer of CAR-T cells can produce remissions in chemo-refractory disease and has changed clinical practice for patients with B cell malignancies^6–10^ and multiple myeloma (MM)^11–13^. Nevertheless, in B cell malignancies more than half of patients do not achieve long-term remissions, and durability of response is still limited in MM. In solid tumors, broad success is lacking^14,15^, and so far, only one αβ-T cell receptor (TCR) based adoptive T cell therapy has obtained regulatory approval in the United States^16^, and not a single CAR-T cell therapy has reached this stage.

To achieve sustained anti-tumor responses in vivo, CAR- or TCR-transgenic T cells must not only recognize and kill tumor cells but also receive distinct co-stimulation and cytokine signals from the environment. In most tumor microenvironments (TMEs) however, co-stimulation is dominated by co-inhibition^17^, and cytokine signals that sustain T stem cell and central memory populations with high proliferative capacity, as well as those supporting effector functions, are lacking or overruled by immunosuppressive signals^18^. Current TCR-based approaches only provide target antigen recognition and completely rely on environmental factors for additional signals. The co-stimulation built into second generation CAR-T cell therapeutics is often insufficient and only partially compensates for the lack of stimulatory environmental signals. So far, most TCR, CAR and co-receptor constructs that equip therapeutic immune cells have been engineered empirically without mechanistic optimization and diversification of their signaling functions, thus limiting the development of more powerful and targeted cellular approaches.

In principle, these limitations could be overcome through the design of biosensing receptors with fully customized molecular properties and associated cellular functions^19,20^. Modern computational protein design techniques can engineer proteins with a wide variety of structures and binding properties^21–23^. However, except for protein-based materials that assemble into specific supramolecular architectures^24^, these methods have mostly been applied to single protein domains^25–27^. The design of dimeric multi-domain signaling receptors remains challenging. The proper orchestration of these receptor functions requires specific ligand-induced protein association, structural switching and long-range communication between domains to ensure signal transduction^28–31^. These properties are essential for the development of biosensors providing precise ligand control of cellular activities but have been neglected so far.

Here, we developed a computational approach for the bottom-up assembly and design of multi-domain receptors with programmable input-output signaling functions. We applied the approach to engineer a new class of receptors that we named T-SenSER (**T**ME-**sen**sing **s**witch receptor for **e**nhanced **r**esponse to tumors). T-SenSERs function as allosteric biosensors and can be co-expressed with conventional CARs in T cells to considerably enhance their function (**Fig. 1a**). T-SenSERs are designed to detect and bind soluble factors in the TME and transmit signals of co-stimulation and common γ-chain cytokines to the engineered T cells. We exploited two distinct soluble factors, vascular endothelial growth factor A (VEGFA) and colony stimulating factor 1 (CSF1), that are enriched in a broad variety of hematologic and solid cancers, respectively. When co-expressed as transgenes with different CARs, T-SenSERs delivered significantly enhanced potency and specificity to CAR-T cells in a TME responsive manner.

**Figure 1.**
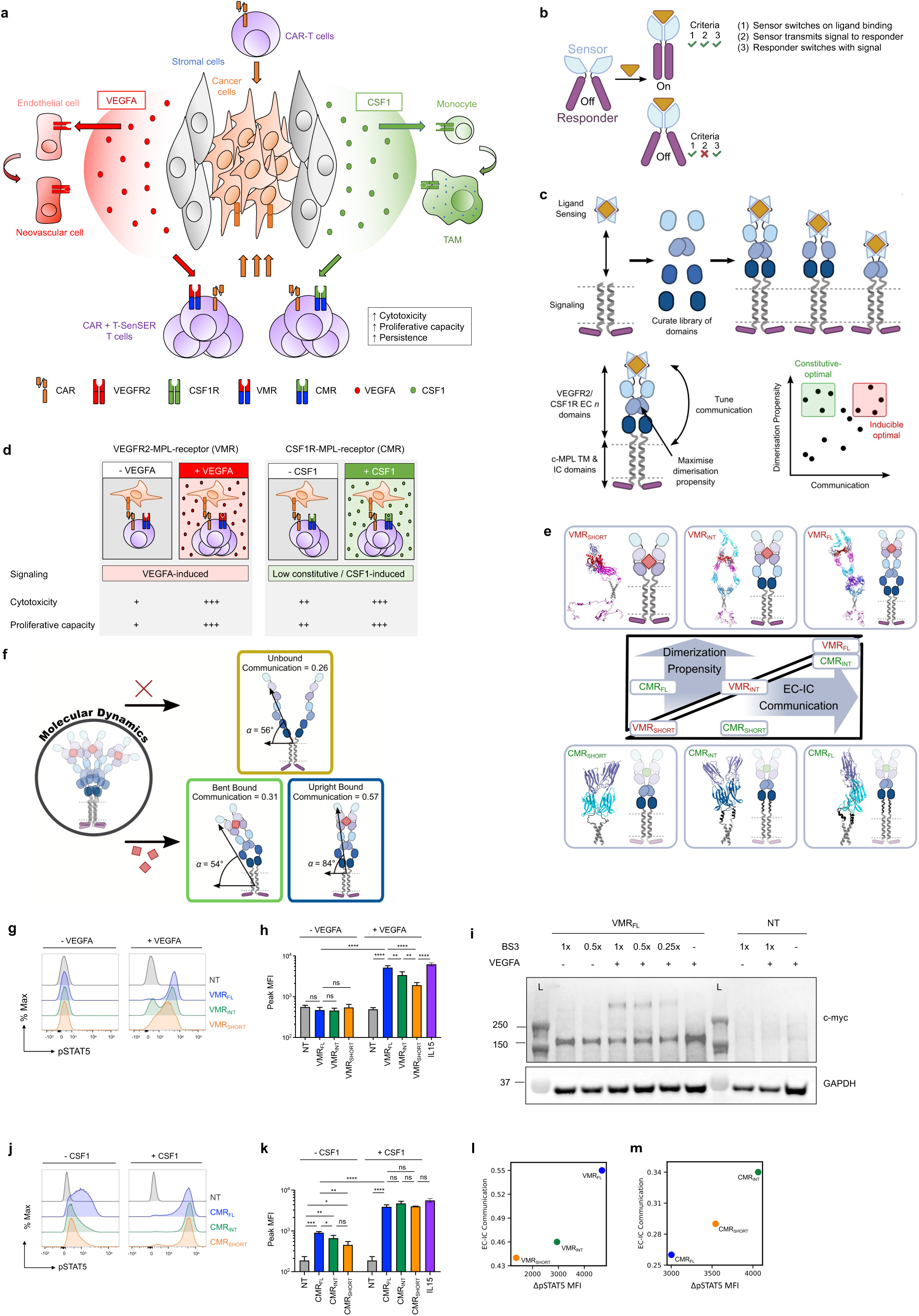
Concept and design of *de novo* assembled biosensors (T-SenSERs). (**a**) Schematic representation of T cells expressing CAR and T-SenSERs responding to vascular endothelial growth factor A (VEGFA) or colony stimulating factor 1 (CSF1) in the tumor microenvironment (TME). CAR: chimeric antigen receptor, T-SenSER: **T**ME **sen**sing **s**witch receptor for **e**nhanced **r**esponse to tumors, TAM: Tumor associated macrophage, VMR: VEGFR2-MPL receptor, CMR: CSF1R-MPL receptor, VEGFR2: vascular endothelial growth factor receptor 2, CSF1R: colony stimulating factor 1 receptor. (**b**) A basic biosensor composed of the sensor and responder elements behaves differently depending on the fulfilled criteria for biosensor activity. (**c**) Overview of the computational platform for the design of de novo assembled T-SenSERs. The desired behavior regimes (constitutive or inducible or low constitutive-inducible) can be met by two scoring metrics, communication and dimerization propensity. (**d**) Concept of inducible VMR T-SenSER activated by VEGFA (left) and low constitutive-inducible CMR T-SenSER responsive to CSF1 (right). VEGFR2-MPL receptor (VMR): Presence of VEGFA induces c-MPL signaling in VMR+ T cells which enhances tumor killing and T cell proliferative capacity. CSF1R-MPL receptor (CMR): Low constitutive c-MPL signaling in CMR+ T cells improves T cell proliferative capacity, homeostatic expansion, and tumor killing. CMR activity is further enhanced with CSF1. (**e**) Schematics and structural models of three different selected T-SenSERs. For VMR: VMR_SHORT_, VMR_INT_ and VMR_FL_. For CMR: CMR_SHORT_, CMR_INT_ and CMR_FL_. Variants were classified by active state dimerization propensity and extracellular-intracellular (EC-IC) communication. (**f**) Selected VMR_FL_ conformations obtained by Molecular Dynamics simulations. An “upright” conformation is observed exclusively for the ligand-bound receptor and enables high communication between sensor and responder. The ligand-free receptor adopts exclusively a “bent” conformation with reduced capacity for communication. Upright and bent conformations likely encode signaling active and inactive states, respectively. (**g**) STAT5 phosphorylation in response to 25 ng/ml VEGFA in human T cells engineered to express VMR_FL,_ VMR_INT_ or VMR_SHORT_ and selected to >96% purity, or non-transduced (NT) controls. Single representative donor FACS histograms, n=5 donors. (**h**) Peak mean fluorescence intensity (MFI) of pSTAT5. n=5 donors, mean±SD, unpaired t-test, 1 representative of 2 independent experiments. Significance levels: *p<0.05, **p<0.01, ***p<0.001, ****p<0.0001, ns: not significant. (**i**) Western Blot for c-myc (tag incorporated in the VMR_FL_ construct) indicating monomeric VMR_FL_ without VEGFA and oligomeric VMR_FL_ with 25ng/ml VEGFA and crosslinking agent bis(sulfosuccinimidyl)suberate (BS3). BS3 dose-dependent detection of VMR_FL_ oligomers. NT T cells served as controls. BS3 concentrations: 1x: 3mM, 0.5x: 1.5mM, 0.25x: 0.75mM. L: ladder, molecular weight indicated on the left, GAPDH: loading control. (**j**) STAT5 phosphorylation in response to 10 ng/ml CSF1 in human T cells engineered to express CMR_FL,_ CMR_INT_ or CMR_SHORT_, or non-transduced (NT) controls. Single representative donor FACS histograms, gated on CMR (CD115) + cells, n=3 donors. (**k**) Peak MFI of pSTAT5. n=3 donors, mean±SD, unpaired t-test. Significance levels: *p<0.05, **p<0.01, ***p<0.001, ****p<0.0001, ns: not significant. (**l, m**) Correlation between the predicted sensor-responder (EC-IC) communication levels of VMR variants (**l**) or CMR variants (**m**) and the measured change in pSTAT5 levels (ΔpSTAT5 MFI: MFI with ligand – MFI without ligand) upon VEGFA (**l**) or CSF1 (**m**) sensing.

## RESULTS

### Computational design of receptors with programmable sensing-response

A membrane receptor biosensor architecture can be schematically decomposed into two elements: 1) an extracellular (EC) ligand binding sensor and 2) an intracellular (IC) signaling responder connected by a transmembrane (TM) domain. Communication between sensor and responder (that we define as coupling below) enables signal transduction across the membrane and activation of intracellular functions upon ligand binding. While the underlying structural mechanisms may vary, sensing-response behaviors rely on at least the following three sequential steps: 1) the sensor changes conformation upon ligand binding; 2) the sensor transmits this structural change to the responder; 3) the responder switches to an active state conformation and triggers receptor activity (**Fig. 1b**).

As in any allosteric system, sensing-response properties can be achieved through diverse design scenarios that impact the receptor’s basal activity, sensitivity, and potency. Specifically, each element can independently switch between an inactive and active state and preferentially occupy one state in isolation. When combined, different biosensor behaviors will be obtained depending on individual bias between inactive/active state and the mechanical coupling between the sensor and responder that will impact the state occupancies. For example, a programmable biosensor scaffold could involve sensor and responder elements that preferentially occupy an inactive and active state conformation, respectively, in isolation and absence of ligand or alternative scenarios. If the sensor and responder are weakly coupled, the responder will readily access the active state and trigger high receptor basal activity (i.e. without ligand). Conversely, strong coupling will maintain the responder mostly in the inactive state, turning off basal activity, while still enabling a strong ligand-induced response.

Studies of natural single pass membrane receptors indicate that while the structural mechanisms underlying biosensing functions can be diverse, they usually involve two main structural modes of activation. In the pre-formed dimer (PFD) mode reported, for example, for the interleukin-7 receptor (IL-7R) or death receptor 5 (DR5) cytokine receptors^32–34^, the receptor self-associates in absence of ligand but mainly occupies inactive state dimer conformations. Ligand binding triggers intra-molecular reorganization propagated allosterically to the cytoplasmic region through coupling, stabilizing active state conformations of the dimer structure. In the monomer-dimer equilibrium shift (MDE) mode initially described for the epidermal growth factor receptor (EGFR)^35^, the ligand-free receptor mostly occupies a monomeric inactive state. Ligand binding triggers receptor association and the formation of active state dimer conformations. However, structural insights into receptor activation are sparse and the prevalence of each mechanism remains highly debated. In fact, several lines of evidence suggest that receptor activation may involve a combination of both allosteric and binding mechanisms^36^.

Since the structural mechanisms of activation of single-pass transmembrane receptors remain largely elusive, we reasoned that a simplified but effective approach for designing receptor biosensors should primarily focus on building optimal active state structures. While this positive design strategy neglects alternative inactive or pre-active states, optimizing structural features for a single target state has proven effective in many protein engineering studies. To account for the diverse activation mechanisms, optimization focuses on the two main structural properties driving signal transduction: dimerization and mechanical coupling, both underlying the MDE and allosteric PFD modes in the active state. In principle, a wide range of constitutive and ligand-inducible activities can be programmed through the modulation of dimerization propensity and communication in the active state. Together, they provide a rational blueprint for engineering receptor scaffolds with diverse and precise sensing and signaling functions (**Figure 1c**)^37,38^.

We developed a computational approach based on these rules to design dimeric biosensors that link the binding of a user-defined chemical input signal to modular cellular responses through genetically encoded fusions of protein domains. Overall, the approach proceeds in the following main steps (**Fig. 1c**): 1) selection of the structural elements defining input, signal transmission and output signals: 1.1) the sensor: extracellular ligand-binding domain and dimerizing domains that link to the responder; 1.2) the responder: transmembrane (TM) and intracellular signaling domains; 2) if necessary, self-association of individual domains into dimeric complexes through docking, and design of juxtamembrane linker sequence connecting the sensor and responder; 3) assembly of multi-domain dimeric scaffolds using structure prediction methods RoseTTAfold and AlphaFold2, and design protocols in Rosetta; 4) ranking of receptor scaffold structures based on their propensity for dimerization and long-range communication (i.e. coupling) between ligand binding and signaling domains that are calculated using Rosetta and Elastic Network models for investigating protein association and coupling, respectively. As mentioned above, this protocol only constructs an ensemble of conformations for the active ligand-bound state of the biosensor. It neglects the impact of the design decisions on alternative states or transitions between states that would require a detailed mechanistic understanding of the activation process. Nevertheless, this positive design selection for optimal oligomerization and coupling in active dimer structures should generate computational libraries of receptors enriched in constructs with desired sensing-response behaviors.

VEGFA and CSF1 were chosen as ligands, two soluble factors that are highly enriched in a variety of TMEs and critically involved in tumor progression. As VEGFs promote neovascularization and CSF1 supports tumor associated macrophage (TAM) development and polarization in various TMEs, these targets open broad fields of applications and offer high translational impact for our T-SenSERs^39,40^. C-MPL signaling was selected as the output signal since we have previously shown that c-MPL activates beneficial costimulatory, cytokine and type I interferon pathways when expressed in TCR-transgenic T cells^41,42^. We aimed to design robust VEGF-MPL-receptor (VMR) scaffolds that are entirely VEGF-dependent, and CSF1-MPL-receptor (CMR) scaffolds that have a low but significant basal activity with full CSF1 ligand-inducibility (‘low constitutive-inducible’). We hypothesized that a low constitutive-inducible CMR has the capacity to counterbalance an immune suppressive TME by enhancing T cell homeostasis and proliferative capacity in the absence of T cell stimulatory cytokines. Thus, low constitutive-inducible CMR activity is expected to sustain local CAR T cell persistence and anti-tumor function in the TME (**Fig. 1d**).

We first analyzed the topology and individual domain structures of the native VEGFR2, CSF1R and c-MPL receptors. While the structures of the full-length receptors remain elusive, several domain structures have been characterized^29,43,44^. Both, VEGFR2 or CSF1R contain immunoglobulin like domains, out of which domains D2 and D3 strongly bind to their cognate ligand. These two domains were selected as the input signaling region of the VMR or CMR respectively. For the output signaling, the structure and activation mechanism of cytokine receptor homologs of c-MPL indicate that strong ligand regulation and potent JAK/STAT signaling is achieved through the intricate coupling between the cytokine TM, juxtamembrane (JM) and the cytoplasmic (CT) regions^45,46^. Hence, we reasoned that an optimal biosensor scaffold should couple the native TM region of c-MPL and not that of VEGFR2 or CSF1R to the c-MPL cytoplasmic domain. In the absence of structural information, we modeled the c-MPL TM domain in a dimer active signaling state from sequence using the method EFDOCK-TM and then assembled the entire signaling (TM+JM+CT) region using our assembly approach. We next curated a library of all known native VEGFR and CSF1R EC domain structures, the recombination of which could modulate coupling between the input and output signaling domains in engineered receptors. Unlike c-MPL, all seven IgG-like VEGFR2 EC domains (D1-7) have been structurally characterized and the isolated D4, D5 and D7 domains are known to homodimerize^29^. Likewise, the D5 and D6 domains are critical for CSF1R homodimerization^44^.

Next, we created a diverse set of chimeras to stringently test our ability to rationally design full-length receptor scaffold structures with fine-tuned signal transduction propensity. Our computational pipeline enables the arbitrary combination of VEGFR or CSF1R EC domains with pre-defined linkers to sample different densities of contacts across the receptor dimerization interface and encode different levels of mechanical coupling. Final designs are returned as a dynamic ensemble of conformations. Thus, the method offers a computationally inexpensive means of obtaining biophysically relevant states from which to assess the modulation of signal transmission triggered by VEGF or CSF1 binding (**Fig. 1c**).

We ultimately designed a total of 18 chimeric receptor scaffolds, with nine constructs generated for each family of sensors. For VMR, these included both intuitive topologies, in which the domain order was preserved as found in natural receptors, as well as non-intuitive designs, where the domain arrangement and combination differed from VEGFR (e.g. VMR_INT_). For CMR, synthetic linkers were designed with properties— such as length, structure, and sequence— that differed from those of CSF1R. We initially explored an extensive space of de novo linker structures and sequences—examining more than 700’000 possibilities—using advanced deep learning methods, including ProteinMPNN and S4PRED, in combination with fragment assembly approaches. From this initial in silico screening, we found that the average helicity of the linkers—critical for dictating coupling properties—was generally low. Our 9 designed CMR chimera combined a diverse range of these ProteinMPNN de novo sequences with a more targeted, structure-informed approach that integrated fragments from native TpoR and CSF1R sequences, leading to linkers with higher helicity and the design of CMR_FL_, CMR_INT_, CMR_SHORT_. Except for CMR_FL_MPNN_helix_, the only chimera from ProteinMPNN to feature helicity, the calculated couplings for the ProteinMPNN linkers fell below the thresholds required to effectively program signaling responses in the CMR constructs.

To conduct in-depth experimental validation, we selected six constructs representative of the dataset and the range of predicted outcomes. For the VMR chimeras, we selected: 1) VMR_FL_, which incorporates all native ectodomains (D1–7), and exhibited the most optimal predicted VEGF response while maintaining the lowest propensity for constitutive activity; 2) VMR_SHORT_, the minimal version of the chimera, where the ligand-binding domain (D1–3) is directly linked to the transmembrane region (TM). It was chosen as a negative control, as it is predicted to exhibit a weak signaling response to VEGF binding; 3) VMR_INT_, a non-intuitive design where D4 is directly connected to D7 (D1–4 + D7), chosen for its intermediate properties. Our calculations predict a gradient of increasing dimerization and coupling properties from VMR_SHORT_ to VMR_INT_ to VMR_FL_, indicating a progressive enhancement in the ability of VMR chimeric scaffolds to redirect VEGF sensing into potent c-MPL signaling (**Fig. 1e**). Among the CMR designs, we selected: 1) CMR_INT_, our top-ranked design in terms of signal transduction propensity; 2) CMR_FL_, which provided a well-balanced compromise between constitutive and ligand-induced activity, aligning with our design goals; 3) CMR_SHORT_, as it exhibited one of the lowest dimerization propensities. Our calculations predicted CMR_FL_ to have the weakest coupling according to our activation model, and therefore should display the highest level of basal activity (**Fig. 1e**). All 3 CMR variants had strong dimerization propensity and were predicted to provide potent signaling responses to CSF1 sensing.

### Molecular Dynamics simulation of VMR_FL_

To validate our design approach and the positive design hypothesis, we next characterized the impact of the ligand on the receptor structure and dynamics using Molecular Dynamics (MD) simulations. Since these calculations are very time-consuming, we selected the most optimal construct in terms of predicted dynamic response, VMR_FL_, and carried out a large-scale simulation both with (total 0.75 μs) and without ligand (1 μs). While these simulations are at least one order of magnitude too short to explore the entire receptor activation process^47^, they revealed distinct conformational properties of the ligand-free and ligand-bound forms that aligned with the expected native behavior of c-MPL^46,48^. PCA and subsequent *K*-means clustering of both the receptor and TM coordinates revealed a unique space occupied by only the ligand-bound state. These conformations corresponded to the receptor adopting an “upright” position, with the representative cluster center displaying an angle of 84° between the lipid membrane and the D2’s center of mass. The remaining conformational spaces shared by both the ligand-bound and ligand-unbound tended towards lower angles of around 55° on average (**Fig.1f**). Calculated mechanical coupling of these representative cluster centers correlated with these angles, with the upright position returning a much higher coupling score than the lower angle ligand-bound or unbound conformations. This finding aligns well with the consensus that inactive receptor tyrosine kinase conformations adopt a bent configuration^49^, and indeed can form direct interactions with the membrane itself^50^, before becoming upright when ligand bound. Overall, our results imply that only conformations accessible to the ligand-bound state can confer the necessary coupling required for a potent response, and that our coupling metric is sensitive enough to capture this behavior.

Consistent with previous experimental findings on c-MPL, the ligand impacts also the conformation of the juxtamembrane (JM) region that flanks the TM on the cytoplasmic side of the membrane, participating in the receptor activation. For instance, a cation-π interaction between R514 and W515 known to stabilize a helical motif in the inactive state^48^ is more often observed in the ligand-free simulations. Conversely, Q516, known to stabilize the interface of the active state^46^, is found with higher frequencies at the interface in the ligand-bound simulation. Overall, while we have not modeled the entire activation process and our simulations have unlikely reached a true inactive state, our ligand-unbound simulations revealed several known inactive-like structural features. The shifts in coupling behaviors and TM-JM motif interactions between the ligand-bound and unbound simulations therefore validate our constructed model and strongly suggest that our assembly protocol can design reasonable receptors with predictable behaviors.

### T-SenSER signal transduction in human T cells

To experimentally explore the computationally predicted signal transduction propensity of the designed VMR and CMR variants, we assessed baseline and VEGFA or CSF1 dependent STAT5 phosphorylation as a surrogate for c-MPL signaling in human T cells transduced with VMR_SHORT_, VMR_INT_, VMR_FL_ or CMR_SHORT_, CMR_INT_, CMR_FL_. We found that all variants were capable of transmitting signal upon VEGFA or CSF1 exposure. For VMR, VMR_FL_ produced the most efficient STAT5 phosphorylation followed by VMR_INT_ and VMR_SHORT_ that were characterized by significantly lower levels of %pSTAT5+ cells and lower peak mean fluorescence intensity (MFI) of pSTAT5 expression when compared to VMR_FL_ or IL15 positive control (**Fig. 1g-h**). Importantly, no spontaneous pathway activation was detected in any of the three VMR variants, indicating that VMRs are fully ligand inducible. To assess whether VEGFA triggers receptor dimerization or intramolecular reorganization of pre-formed dimers, we characterized the size of VMR_FL_ in transgenic T cells by Western Blot in presence or absence of VEGFA. To facilitate detection of receptor oligomers, samples were treated with the crosslinking reagent BS3 (**Methods**). In the absence of VEGFA, we found VMR_FL_ as monomers, while in the presence of VEGFA and BS3, VMR_FL_ was detectable as ligand-bound oligomers (**Fig. 1i**). These observations imply that VMR_FL_ signaling is at least partly controlled by VEGFA-induced dimerization. For CMR, CMR_FL_ had the strongest constitutive baseline activity in the absence of CSF1, while CMR_INT_ and CMR_SHORT_ had significantly lower baseline activity. In addition, all CMR variants were highly inducible in the presence of the ligand CSF1 (**Fig. 1j-k**).

Overall, the measured signal transductions are consistent with the intended designed properties. Higher coupling in VMRs locks the responder in the inactive monomeric state in absence of ligand, hence lowering constitutive activity while promoting potent switching and activation upon ligand binding (**Fig. 1l**). The stronger coupling in VMR_FL_ results in ligand binding driving a higher proportion of receptors into the active state than in VMR_INT_ and VMR_SHORT_. Due to lower communication, the c-MPL responder in CMRs often occupies the active state and triggers constitutive activity (**Fig. 1m**). The lower communication in CMR_FL_ results in higher basal activity than in CMR_INT_ and CMR_SHORT_ while maintaining maximal activity in presence of CSF1. Additionally, within each family of sensors, we observed a linear relationship between the ligand-induced shifts in activity (i.e. induced – constitutive) and the calculated coupling values (**Fig. 1l-m**). These findings indicate that our calculated coupling metrics are strong determinants of receptor signaling activity.

### T-SenSER pathway activation and signaling thresholds in vitro and in vivo

The first step in c-MPL signaling is activation of the JAK/STAT pathway leading to phosphorylation of STAT5 and STAT3. In addition, c-MPL also activates the PI3K/AKT/mTOR axis as well as the MAPK/ERK1/2 pathway (**Fig. 2a**)^51^. To gain deeper insight into pathway activation upon VMR_FL_, CMR_FL_ or c-MPL activation in transgenic human T cells by the respective recombinant ligands (VEGFA, CSF1, thrombopoietin (TPO)), we assessed phosphorylation of STAT5, STAT3, S6, ERK1/2 and p38 MAPK. We found that the c-MPL endodomain used in VMR_FL_ and CMR_FL_ reliably transmitted signals activating all evaluated components, and that the profile of ligand dependent pathway activation was comparable between VMR_FL_, CMR_FL_ and c-MPL transgenic T cells (**Figs. 2b-f**).

**Figure 2.**
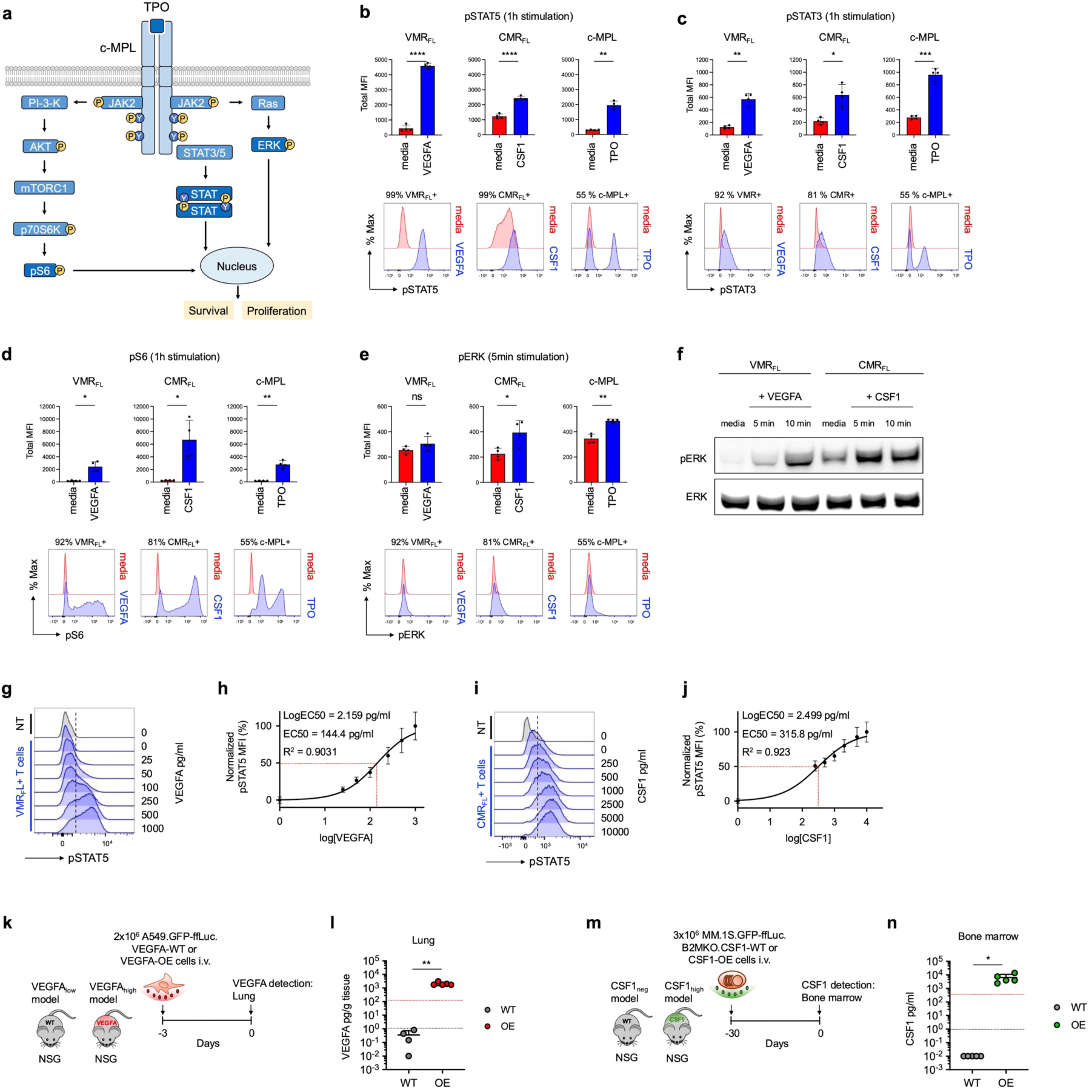
Characterization of T-SenSER phospho-signaling and ligand sensitivity. (**a**) Schematic of the c-MPL signaling pathway. (**b-e**) Total mean fluorescence intensity (MFI) of VMR_FL_, CMR_FL_ or c-MPL engineered T cells for (**b**) pSTAT5, (**c**) pSTAT3, (**d**) pS6, (**e**) pERK in response to 25ng/ml VEGFA (for VMR_FL_), 10ng/ml CSF1 (for CMR_FL_) or 25ng/ml TPO (for c-MPL) (top) and representative donor FACS histograms (bottom). The purity of the analyzed population is indicated above the FACS histograms. n=4 donors, mean±SD, unpaired t-test with Welch’s correction. Significance levels: *p<0.05, **p<0.01, ***p<0.001, ****p<0.0001, ns: not significant. (**f**) Western Blot of pERK and total ERK in VMR_FL_ or CMR_FL_ engineered T cells and baseline (media) and upon incubation for 5 or 10 min with 25ng/ml VEGFA or 10ng/ml CSF1, respectively. (**g**) VEGFA dose-dependent STAT5 phosphorylation in VMR_FL_+ T cells on >96% VMR+ purified cells. FACS histograms of a representative donor of n=5 donors. (**h**) Non-linear curve fit and calculation of EC50 (red dotted line) for pSTAT5 MFI. n=5 donors, mean±SD. (**i**) CSF1 dose-dependent STAT5 phosphorylation in CMR_FL_+ T cells on CD115+ gated cells. FACS histograms of a representative donor of n=3 donors. (**j**) Non-linear curve fit and calculation of EC50 (red dotted line) for pSTAT5 MFI. n=3 donors, mean±SD. Measurement of NT T cells was used for the origin data point. (**k**) Schematic of mouse models to determine VEGFA levels in mice injected intravenously (i.v.) with A549.GFP-ffLuc.VEGFA-WT or VEGFA-OE cells. (**l**) VEGFA levels in lung tumor tissue lysates of NSG mice 3 days after tumor cell injection, n=4-5 mice/group, mean±SD, unpaired t-test with Welch’s correction. (**m**) Schematic of mouse models to determine CSF1 levels in mice injected intravenously (i.v.) with MM.1S.GFP-ffLuc.B2MKO.CSF1-WT or CSF1-OE cells. (**n**) CSF1 levels in bone marrow lysates of NSG mice 30 days after tumor cell injection, n=5 mice/group, mean±SD, unpaired t-test with Welch’s correction. (**l**, **n**) Significance levels: *p<0.05, **p<0.01. EC50 (red dotted line) and detection threshold, linear range (black dotted line).

Both serum VEGF and CSF1 levels have been intensively studied across tumor histologies and are significantly higher in cancer patients than in healthy individuals^52,53^. Body compartment distribution of VEGFs and CSF1 is altered in cancer, indicating that T-SenSER T cells will likely encounter higher VEGF or CSF1 levels in malignant than in normal tissues, thereby mediating enhanced tumor specificity. To determine the activation threshold and EC50 of both VMR_FL_ and CMR_FL_ in response to ligand, we quantified pSTAT5 levels in VMR_FL_+ and CMR_FL_+ T cells in response to increasing concentrations of VEGFA or CSF1 (**Fig. 2g-j**). EC50 for VMR_FL_ was 144 pg/ml (**Fig.2h**) while the EC50 for CMR_FL_ was 315.8 pg/ml (**Fig.2j**).

To evaluate our strategy in different cancer models, we selected metastatic lung cancer and multiple myeloma (MM). Metastatic lung cancer is a solid tumor that is difficult to cure despite the introduction of immune checkpoint blockade therapy^54^, and CAR-T or TCR-T cells are still in early development. MM has approved CAR T cell therapies, but current products and indications do not confer long-lasting remissions^12^, thus enhanced targeting moieties or combinations with novel approaches are highly warranted. To model the various VEGFA or CSF1 levels in mouse xenografts and compensate for the lack of a human TME as a source of human VEGFA and CSF1, we engineered A549.GFP-ffLuc cells with VEGFA and MM.1S.GFP-ffLuc cells with CSF1 overexpression (OE). In vitro, A549.GFP-ffLuc.VEGFA-wild-type (WT) cells produced low levels of VEGFA, contrary to A549.GFP-ffLuc.VEGFA-OE cells which secreted VEGFA levels above the VMR_FL_ EC50 threshold. Next, A549 cells were engrafted intravenously in immunocompromised NOD-SCID-γ-chain^-/-^ (NSG) mice and lung tissue levels of VEGFA were determined after three days (**Fig. 2k-l**). VEGFA levels in the lung remained below the EC50 for VMR_FL_ activation in A549.VEGFA-WT engrafted mice (VEGFA_low_ model), while a median of 1.775 ng VEGFA/g total protein (range 1.725 – 3.061 ng/g, n=5) was reached in lung tissues of A549.VEGFA-OE engrafted mice, significantly above the EC50 for VMR_FL_ activation (VEGF_high_ model) (**Fig. 2l**). VEGFA tissue levels in the VEGF_high_ model was on average 126-fold lower than the levels reported in lung cancer patients (median 224 ng VEGFA/g total protein, range 30-1870, n=71)^55^. The VEGF_high_ model is therefore appropriate to evaluate CAR.VMR_FL_+ T cell function in vivo but may underestimate VMR_FL_ potency due to lower tissue levels in the animal model compared to patient tissues. In MM, MM.1S.GFP-ffLuc.B2MKO.CSF1-WT cells did not produce detectable CSF1 in vitro. However, high levels of human CSF1 were detected from engineered MM.1S.GFP-ffLuc.B2MKO.CSF1-OE cells, above the CMR_FL_ EC50 levels. To quantify CSF1 levels in vivo, we engrafted both types of MM.1S cells intravenously in NSG mice and analyzed BM lysates (**Fig. 2m-n**). In BM of mice engrafted with MM.1S.GFP-ffLuc.B2MKO.CSF1-OE, we detected CSF1 levels above the CMR_FL_ EC50 with a median of 3.998 ng/ml (range 2.360 – 13.878 ng/ml, n=5), while no CSF1 was detected in BM of MM.1S.GFP-ffLuc.B2MKO.CSF1-WT engrafted mice (**Fig. 2n**). We termed our two models CSF1_high_ and CSF1_neg_ model respectively and used these to characterize CMR_FL_+ CAR-T cell function in vivo.

To maximize the impact of the designed T-SenSERs on CAR T-cell functions, we selected VMR_FL_, the VMR construct with the highest signaling response to VEGF, and CMR_FL_, the CMR construct combining the strong response to CSF1 with the highest basal activity to favor also constitutive homeostasis and enhanced effector function in absence of cytokines. To evaluate T-SenSER activity in vivo in relation to the levels of ligand present in the TME we established animal models with different levels of VEGFA or CSF1 that reflect the clinical situation of lung cancer and myeloma patients.

### Consequences of VMR_FL_ activation in human CAR-T cells targeting lung cancer in vitro

VMR_FL_ was efficiently co-transduced in activated human T cells with conventional second generation (28ζ, BBζ) or non-signaling control (Δ) CARs targeting the antigen Ephrin A2 (EphA2). CAR expression levels were comparable between CAR and CAR.VMR_FL_ transduced cells, and VEGFA-induced STAT5 phosphorylation was comparable between VMR_FL_ and CAR.VMR_FL_+ T cells (**Fig. 3a-b**)^56^. The 4H5 single chain variable fragment recognizes a conformational epitope of EphA2 that is exposed on a wide variety of malignant but not on normal cells, including A549 lung cancer cells^57^. The impact of VMR_FL_ activation on tumor killing and T cell expansion was assessed in sequential co-cultures where T cells were repetitively challenged with fresh tumor cells ± VEGFA (**Fig. 3c-d**). Tumor killing and T cell expansion were quantified after each challenge. Full T cell activation with target cell killing and sustained T cell expansion occurred in the presence of both tumor cells and VEGF (**Fig. 3e-g**). Cytotoxicity was sustained in vitro even in T cells transduced with CAR alone (**Fig. 3e**), but T cell expansion was enhanced in the presence of VEGFA and VMR_FL_ signaling (**Fig. 3f-g**). A slight but not significant enhancement of both killing and T cell expansion with CAR.VMR_FL_+ T cells was observed without exogenous VEGFA addition. This is probably due to low levels of VEGFA production by A549-WT cells used in the assay (**Fig. 3e-g**). VMR_FL_ activity in the presence of VEGFA did not alter cytokine nor cytolytic granule production when compared to CAR alone and did not impact the T cell subset composition and differentiation status of T cells.

**Figure 3.**
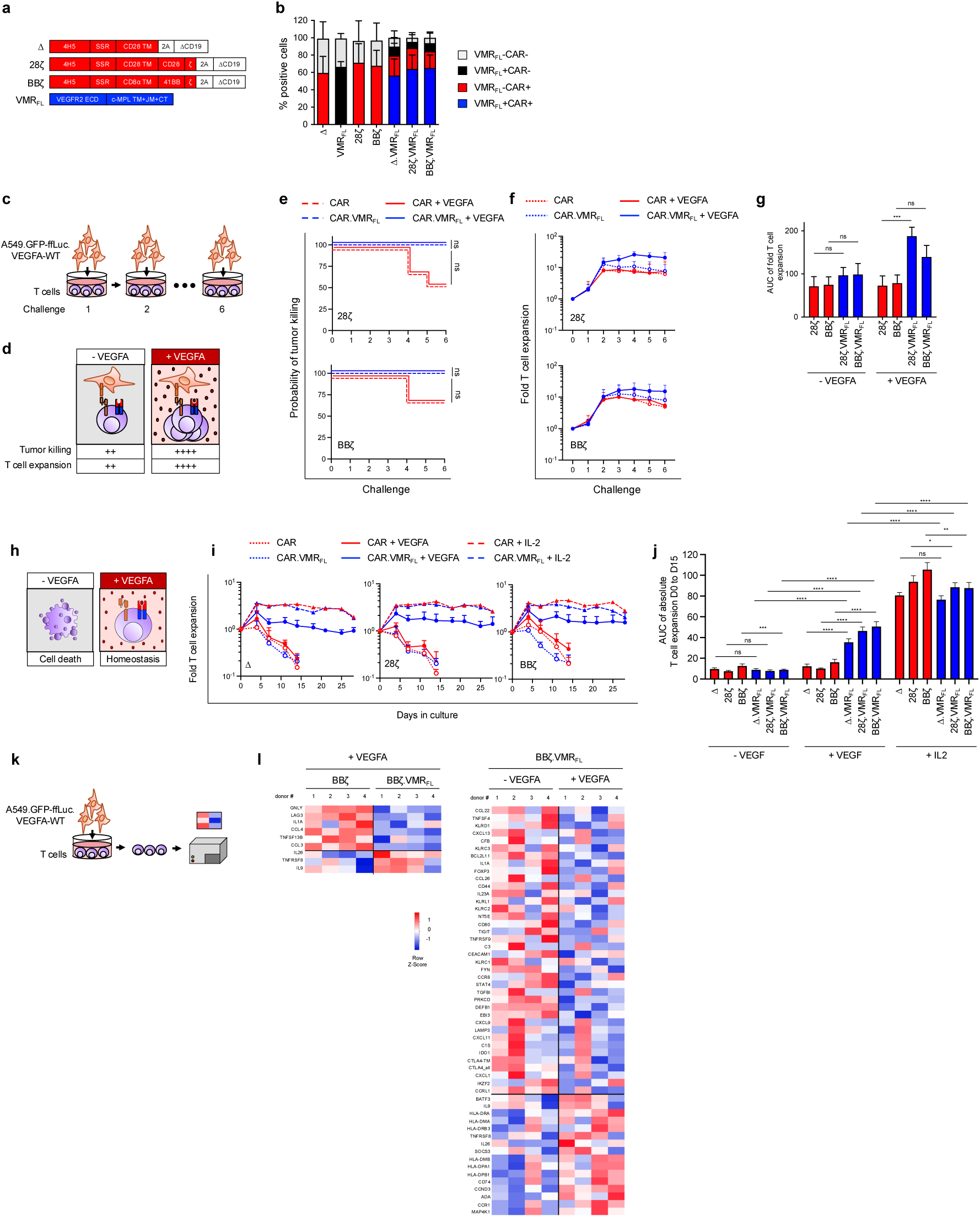
Effects of VEGFA-dependent VMR_FL_ activation on EphA2-CAR T cell cytotoxicity, expansion and transcriptional profile targeting lung cancer. (**a**) Schematic representation of the retroviral vector constructs. 4H5: Single-chain variable fragment targeting ephrin type-A receptor 2 (EphA2). Δ: Non-signaling control CAR. 28ζ: Second generation CAR with CD28 and CD3ζ endo-domains. BBζ: second generation CAR with 41BB and CD3ζ endo-domains. ECD: extracellular domain. TM: Transmembrane region. JM: Juxtamembrane region. CT: Cytoplasmic region. SSR: Short spacer region. (**b**) Co-transduction efficiencies of activated T cells by FACS for each construct or combinations thereof. CD19 staining: marker of CAR transduction, VEGFR2 staining: direct detection of VMR_FL_, n=8 donors (n=6 for Δ and Δ.VMR_FL_ conditions), mean±SD. % positive cells and transgene detection are color coded. (**c**) Schematic of the sequential co-culture assay with repetitive tumor challenge every 3-4 days. (**d**) Schematic of co-culture conditions and expected results. (**e**) Probability of tumor killing in the sequential co-culture assay ± exogenous VEGFA (25 ng/ml, n=7 donors), Kaplan Meier analysis with log-rank (Mantel-Cox) test. (**f**) Fold T cell expansion in sequential co-cultures, n=7 donors, mean±SD. (**g**) Area under the curve (AUC) analysis of fold T cell expansion shown in panel **f**, from challenge 1 to 6, mean±SEM, unpaired t-test with Welch’s correction. (**h**) Schematic of T cell culture conditions in the absence of tumor and expected results. (**i**) T cell expansion and survival over time in media ± VEGFA 25 ng/ml or IL2 50U/ml with no tumor challenge. n=4 donors, mean±SD. (**j**) Area under the curve analysis of absolute T cell expansion shown in panel **i**, from day 0 to day 15, mean±SEM, unpaired t-test with Welch’s correction. (**k**) Schematic of experiment for differential gene expression analysis. (**l**) Differential gene expression analysis by Nanostring. BBζ+ or BBζ.VMR_FL_+ T cells were purified after 1 tumor challenge with VEGFA (25ng/ml) (left) and BBζ.VMR_FL_+ T cells were isolated after 1 tumor challenge ± VEGFA (25ng/ml) (right). Significantly upregulated (red) or downregulated (blue) genes are shown. n=4 donors. (**e, g, j**) Significance levels: *p<0.05, **p<0.01, ***p<0.001, ****p<0.0001, ns: not significant.

To assess the potential for long-term persistence of engineered T cells in the absence of EphA2+ tumor, we performed a 4-week homeostatic maintenance experiment. VMR_FL_ activation by VEGFA alone provided T cell homeostasis and survival. In line with our previous observations on transgenic c-MPL signalling in human T cells^42^, overall expansion levels were significantly increased with VEGFA compared to media alone but remained below those of IL-2 control. (**Fig. 3h-j**).

Lastly, we hypothesized that VMR_FL_ signaling is complementary to EphA2-CAR BBζ signaling since the pathways activated by the endo-domains are largely distinct. Thus, we assessed differential gene expression analyzing global immune response signatures in BBζ.VMR_FL_+ T cells after one in vitro tumor challenge ± VEGFA, and also compared BBζ with BBζ.VMR_FL_+ T cells in the presence of VEGFA. We identified several highly differentially expressed genes that are associated with enhanced T cell co-stimulation (e.g. CD80, TNFRSF8, HLA class II molecules) or enhanced effector function (e.g. GNLY) in cells with VMR_FL_ stimulation. VMR_FL_ activation also led to a reduction in expression of genes associated with T cell exhaustion (e.g. CTLA4, LAG3, TIGIT), or factors associated with immune suppression (e.g. reduced transcription of NT5E, TGFB1, increased transcription of ADA) (**Figs. 3k-l**).

These results suggest that, as intended, VMR_FL_ activation delivered signals for CAR-T cell expansion and persistence, and involved transcriptional changes associated with co-stimulation, effector function, and reduced exhaustion in combination with a BBζ CAR.

### Consequences of CMR_FL_ activation in human CAR-T cells targeting multiple myeloma in vitro

CMR_FL_ was co-expressed efficiently in activated human T cells with A Proliferation inducing ligand (APRIL) based CARs targeting two antigens expressed on MM: B cell maturation antigen (BCMA) and transmembrane activator and CAML interactor (TACI). Monomers of APRIL (m) were used as ligand binding domains and CARs were expressed in conventional first (mζ) or second generation (m28ζ, mBBζ) format as previously described (**Fig. 4a-b**)^58^. CAR cell surface expression levels were slightly lower in CAR.CMR_FL_ compared to CAR T cells, STAT5 phosphorylation levels were slightly lower in CAR.CMR_FL_ compared to CMR_FL_ T cells. The impact of CMR_FL_ activation on tumor killing and T cell expansion was assessed in sequential co-cultures (**Fig. 4c-d**) with two different MM cell lines expressing different target antigen levels (NCI-H929 (BCMA++TACI-) and MM.1S (BCMA+TACI+)). CMR_FL_ expression provided a significant advantage for sequential killing of NCI-H929 cells only in combination with the mBBζ CAR, while killing of MM.1S cells was enhanced with all three evaluated mAPRIL-based CARs (mζ, m28ζ, mBBζ). The CMR_FL_ constitutive baseline activity was sufficient to enhance killing, which was not further improved with the addition of CSF1 at the tested E:T ratio (**Fig. 4e**). CMR_FL_ also boosted T cell expansion in vitro that was significantly higher in all conditions except with the m28ζ CAR targeting NCI-H929 (**Fig. 4f-g**). CMR_FL_ activity did not alter cytokine nor granzyme production when compared to CAR alone. During sequential co-culture, the CD4/CD8 ratio changed with an enrichment in CD8+ T cells. The effector/ memory differentiation status was mostly dictated by the type of CAR and not significantly impacted by CMR_FL_ activity. To assess the impact of CMR_FL_ activity on T cell exhaustion/ activation phenotype, we assessed expression of LAG3, CTLA4, TIGIT, PD1 and TIM3 at baseline and after 5 tumor challenges and compared % positive populations and marker MFI on CAR+ and CAR.CMR_FL_+ T cells. We found a trend to decreased expression of LAG3, CTLA4 and PD1 on mBBζ CAR.CMR_FL_+ CD4+ T cells, and increased TIM3 on mBBζ CAR.CMR_FL_+ CD8+ T cells. Lastly, in the 4-week homeostatic maintenance experiment in the absence of tumor, we found that the CMR_FL_ constitutive baseline activity in T cells was sufficient to mediate significant low-level expansion and survival. Addition of CSF1 significantly enhanced expansion and survival of CMR_FL_+ T cells, comparable to the levels of NT controls supplemented with IL2. Adding IL2 to CMR_FL_+ T cells significantly augmented the effects of the baseline constitutive activity, indicating that the combination of c-MPL signaling and native common γ-chain cytokine signals mediated by IL2 are complementary (**Figs. 4h-j**).

**Figure 4.**
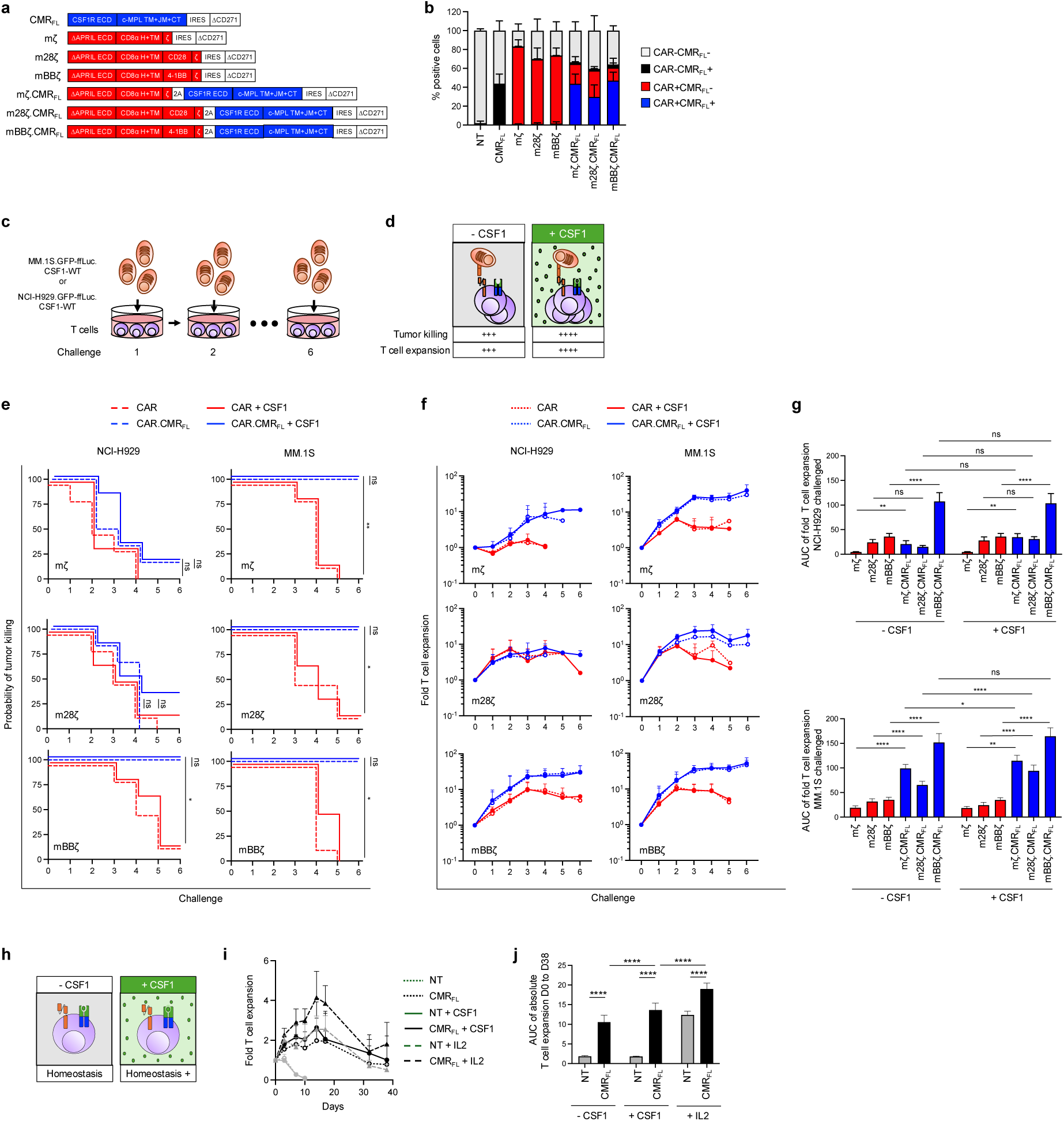
Effects of constitutive baseline and CSF1-dependent CMR_FL_ activation on mAPRIL CAR-T cell cytotoxicity and expansion targeting multiple myeloma. (**a**) Schematic representation of the retroviral vector constructs. ΔCD271: Truncated nerve growth factor receptor as marker for transduction. ΔAPRIL ECD: Truncated monomeric APRIL extracellular domain. mζ: First generation CAR with CD3ζ endo-domain. m28ζ: Second generation CAR with CD28 and CD3ζ endo-domains. mBBζ: Second generation CAR with 41BB and CD3ζ endo-domains. mζ.CMR_FL_, m28ζ.CMR_FL_ and mBBζ.CMR_FL_: Polycistronic vectors with CARs and CMR_FL_ separated by a 2A sequence. (**b**) Transduction efficiencies of activated T cells by FACS for each construct. CD271 staining: marker of CAR or CAR.CMR_FL_ transduction, CD115 (CSF1R) staining: direct detection of CMR_FL_, n=6 donors (n=3 for m28ζ.CMR_FL_), mean±SD. % positive cells and transgene detection are color coded. (**c**) Schematic of the sequential co-culture assay with repetitive tumor challenge every 3-4 days. (**d**) Schematic of co-culture conditions and expected results. (**e**) Probability of tumor killing in the sequential co-culture assay ± CSF1 (10 ng/ml), n=6 donors, (n=3 for mζ.CMR_FL_ and m28ζ.CMR_FL_ challenged with MM.1S). Kaplan-Meier analysis with log-rank (Mantel-Cox) test. (**f**) Fold T cell expansion in sequential co-cultures. n=6, (n=3 for mζ.CMR_FL_ and m28ζ.CMR_FL_ challenged with MM.1S) mean±SD. (**g**) Area under the curve (AUC) analysis of fold T cell expansion shown in panel **f**, from challenge 1 to 6, mean±SEM, unpaired t-test with Welch’s correction. (**h**) Schematic of T cell culture conditions in the absence of tumor and expected results. (**i**) T cell expansion and survival over time in media ± CSF1 10 ng/ml or IL2 50U/ml, with no tumor challenge. n=5, mean±SD. (**j**) Area under the curve (AUC) analysis of absolute T cell expansion shown in panel **i**, from day 0 to day 38, mean±SEM, unpaired t-test with Welch’s correction. (**e, g, j**) Significance levels: *p<0.05, **p<0.01, ***p<0.001, ****p<0.0001, ns: not significant.

These results demonstrate that the constitutive baseline activity of CMR_FL_ enhanced the homeostatic expansion capacity of engineered T cells and increased the in vitro sequential killing and expansion capacity of mAPRIL CAR T cells in situations of high tumor loads.

### In vivo potency and selectivity of VMR_FL_+ EphA2-BBζ CAR T cells in lung cancer

We next evaluated the in vivo impact and VEGF-dependence of VMR_FL_ function in EphA2-BBζ CAR T cells. In the systemic VEGF_low_ model, NSG mice engrafted with A549.GFP-ffLuc.VEGFA-WT cells were treated with a single limiting dose of 1×10^5^ T cells. Bioluminescent imaging (BLI) revealed partial response to BBζ CAR T cell therapy, and as expected no impact of VMR_FL_ addition was detected. In the systemic VEGF_high_ model, mice were engrafted with A549.GFP-ffLuc.VEGFA-OE cells. Mice treated with Δ or BBζ CAR T cells had rapidly progressive disease and reached the experimental endpoint within ten days. In contrast, mice treated with BBζ.VMR_FL_+ T cells mounted a potent anti-tumor response, associated with a significant reduction in VEGFA serum levels. Most importantly, the overall survival of mice treated with BBζ.VMR_FL_+ T cells was significantly enhanced compared to BBζ CAR-T cell treated mice (**Fig. 5a-e**).

**Figure 5.**
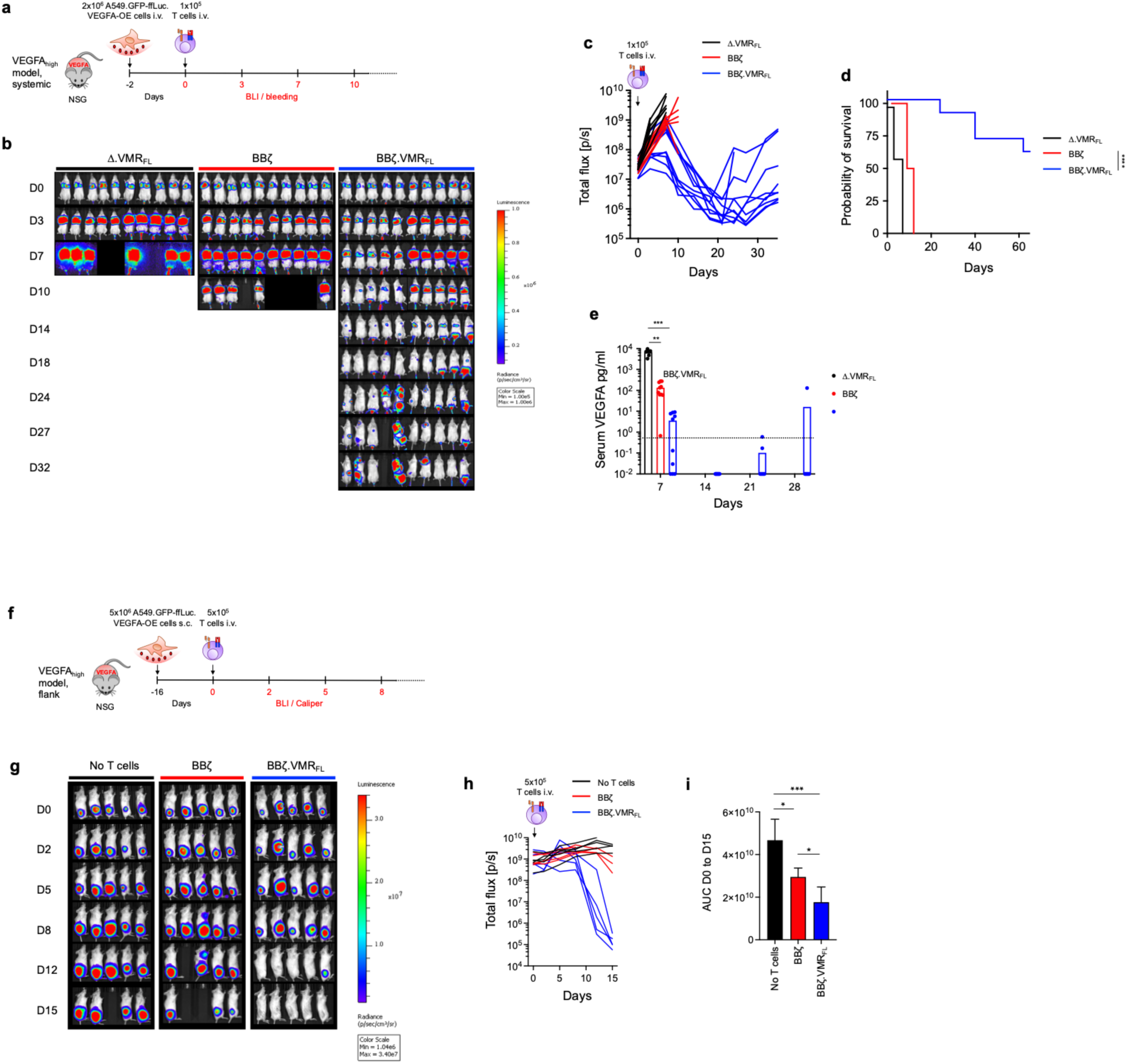
VEGFA-dependent in vivo anti-tumor function of EphA2-BBζ.VMR_FL_ T cells in lung cancer. (**a**) Schematic of the VEGFA_high_ systemic mouse model. (**b**) Individual mouse images, color scale ranges from 1×10^5^ to 1×10^6^ p/sec/cm2/sr. (**c**) Summary of total flux, lines representing values from individual mice. n=10 mice/group, pooled results of 2 independent experiments. (**d**) Survival of mice. n=10 mice/group, Kaplan Meier analysis, log-rank (Mantel-Cox) test. (**e**) Human VEGFA levels in serum of surviving mice. Black dotted line: detection threshold, linear range. Unpaired t-test with Welch’s correction. (**f**) Schematic of the VEGFA_high_ subcutaneous mouse model. (**g**) Individual mouse images, color scale ranges from 1.04×10^6^ to 3.40×10^7^ p/sec/cm2/sr, n=5 mice/group. (**h**) Summary of total flux, lines representing values from individual mice, n=5 mice/group, 1 representative of 2 experiments. (**i**) Area under the curve analysis of total flux shown in panel **h** from day 0 to day 15, mean±SEM, unpaired t-test with Welch’s correction. (**d, e, i**) Significance levels: *p<0.05, **p<0.01, ***p<0.001, ****p<0.0001, ns: not significant.

To evaluate whether VMR_FL_ also enhanced BBζ CAR T cell function in a subcutaneous solid tumor model, we engrafted A549.GFP-ffLuc.VEGFA-OE cells subcutaneously in the flanks of NSG mice. After stable tumor engraftment, mice were treated with a limiting dose of 5×10^5^ T cells intravenously and anti-tumor activity was assessed by BLI. We found again a significantly better anti-tumor response in mice treated with BBζ.VMR_FL_+ T cells with clearance of their tumors within 2 weeks, compared to BBζ CAR T cells or untreated controls (**Fig. 5f-i**), confirming the results obtained with the systemic VEGF_high_ model.

Thus, EphA2-BBζ CAR-T cells equipped with the VMR_FL_ T-SenSER provided potent VEGFA-dependent in vivo anti-tumor activity in both systemic and subcutaneous lung cancer xenograft models and favored tumor eradication.

### In vivo potency and selectivity of CMR_FL_+ BBζ CAR T cells in multiple myeloma

Lastly, we assessed both the low constitutive and CSF1-dependent CMR_FL_ mediated enhancement of CAR-T cells targeting MM. In the CSF1_neg_ model, MM.1S.GFP-ffLuc.B2MKO.CSF1-WT engrafted NSG mice were treated with a single dose of 5×10^6^ T cells using the mAPRIL CAR as model system. Significant but transient anti-tumor activity was observed in the mBBζ but also in the mBBζ.CMR_FL_ treatment groups, with no benefit observed by the co-expression of CMR_FL_. Thus, unlike the in vitro results, the constitutive baseline activity of CMR_FL_ was not sufficient to enhance mAPRIL CAR T cell potency in vivo. To test whether endogenous tissue levels of human CSF1 are sufficient to mediate enhanced anti-tumor activity to CAR.CMR_FL_ T cells, we used NSG-Quad mice that express transgenic human CSF1 in tissues as recipients (CSF1_endo_ model).^59^ We systemically engrafted MM.1S.GFP-ffLuc.B2MKO.CSF1-WT cells, treated mice with fully human heavy-chain-only BCMA directed FHVH33-BBζ or FHVH33-BBζ.CMR_FL_ T cells, and followed tumor growth by BLI (**Fig. 6a**).^60^ We found more potent anti-tumor responses with FHVH33-BBζ.CMR_FL_ than with FHVH33-BBζ T cells at limiting dose. Tumor progression was significantly delayed, and survival of mice prolonged in the FHVH33-BBζ.CMR_FL_ treatment group. Finally, in the CSF1_high_ model, co-expression of CMR_FL_ along with a CAR also significantly improved outcomes of mice, either when combined with the mBBζ or the FHVH33-BBζ CAR (**Fig. 6f-l**). We found significantly enhanced tumor control (**Fig. 6h-j**) and improved survival of mice (**Fig. 6k-l**) at limiting T cell doses.

**Figure 6.**
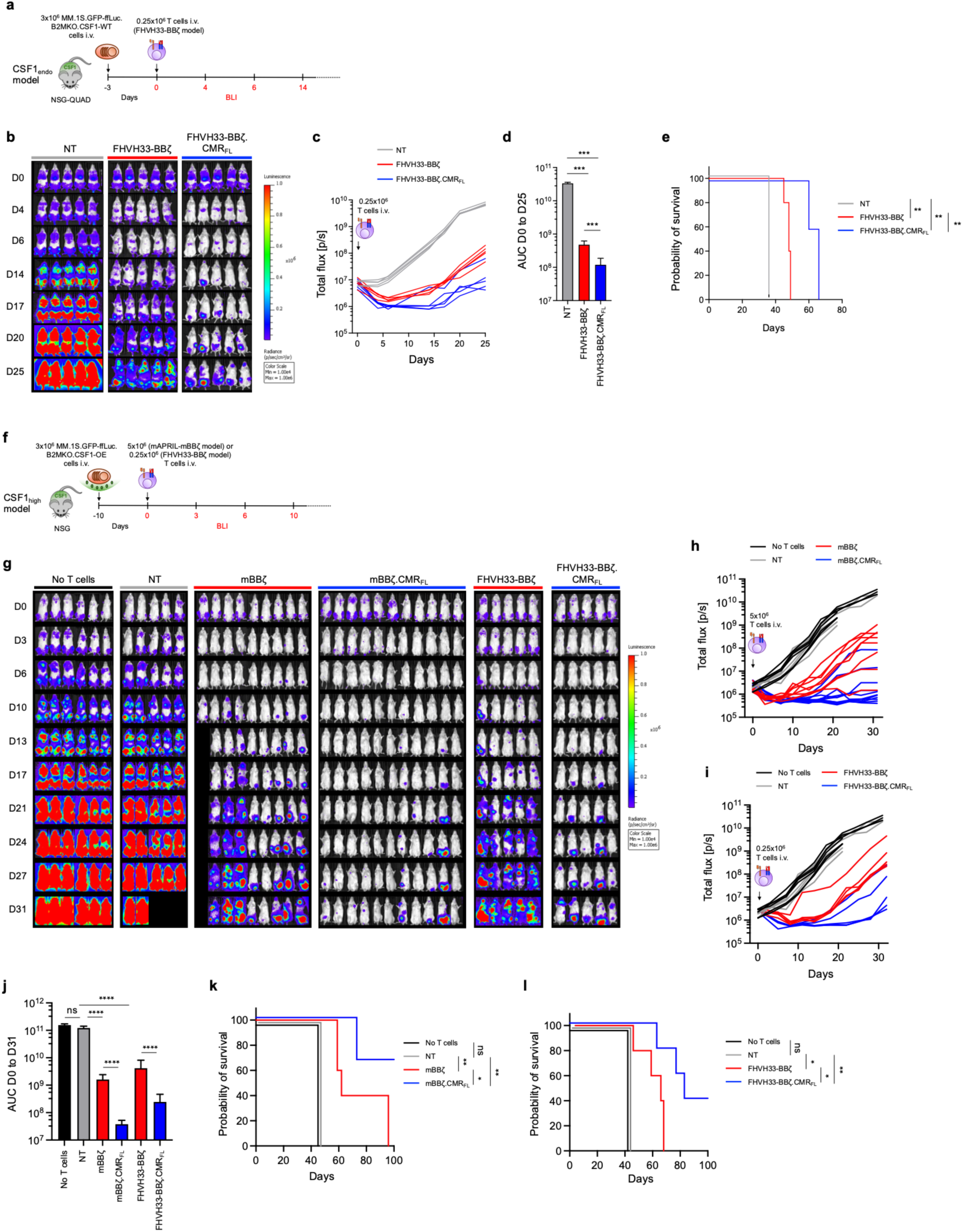
CSF1-dependent in vivo anti-tumor function of APRIL-mBBζ.CMR_FL_ and FHVH33-BBζ.CMR_FL_ T cells in multiple myeloma. (**a**) Schematic of the CSF1_endo_ mouse model evaluating the in vivo anti-tumor function of engineered T cells in NSG-Quad mice. (**b**) BLI images of individual mice, color scale ranges from 1×10^4^ to 1×10^6^ p/sec/cm2/sr. (**c**) Summary of total flux, lines representing individual mice. n=5 mice/group, results of 1 experiment. (**d**) Area under the curve analysis of total flux shown in panel **c**, from day 0 to day 25 mean±SEM, unpaired t-test with Welch’s correction. (**e**) Survival of mice. Kaplan Meier analysis, log-rank (Mantel Cox) test. (**f**) Schematic of the CSF1_high_ mouse model. (**g**) Images of individual mice, color scale ranges from 1×10^4^ to 1×10^6^ p/sec/cm2/sr. (**h, i**) Summary of total flux, lines representing individual mice. n=5 mice/group (except for n=6 for untreated, n=9 for mBBζ, n=11 for mBBζ.CMR_FL_ groups), pooled results of 2 independent experiments for mAPRIL CAR model, 1 experiment for FHVH33 CAR model. (**j**) Area under the curve analysis of total flux shown in panel **h** and **i**, from day 0 to day 31, mean±SEM, unpaired t-test with Welch’s correction. (**k, l**) Survival of mice. n=5 mice/group from 1 experiment, Kaplan Meier analysis, log-rank (Mantel Cox) test. (**d, e, j, k, l**) Significance levels: *p<0.05, **p<0.01, ***p<0.001, ****p<0.0001, ns: not significant.

Thus, our results demonstrate that the CMR_FL_ T-SenSER significantly enhanced anti-tumor activity of CAR-T cells against MM in a CSF1-dependent manner. Enhanced T cell potency was achieved both in response to physiological tissue levels of human CSF1 in NSG-Quad mice and in an engineered cell line model with CSF1 overexpression by tumor cells.

## Discussion

We developed a computational method for the bottom-up assembly and design of receptors with programmable signaling activity. We applied our method to create T-SenSERs with predictable signaling responses to soluble factors enriched in TMEs and validated our assembly protocol through large-scale Molecular Dynamics simulations. T-SenSER’s constitutive and ligand-induced signaling were computationally tuned to enhance the context-dependent potency of CAR-T cells. We demonstrate that both VEGFA and CSF1, enriched in TMEs of various tumor types, can be exploited as chemical cues for activating synthetic signaling through VMR_FL_ or CMR_FL_ expressed in CAR-T cells, respectively, and overcome the lack of endogenous co-stimulation and cytokine signaling in TMEs. Both VMR_FL_ and CMR_FL_ increased the potency of limiting doses of CAR T cells in ligand-rich lung cancer and MM xenograft models. Our approach has the potential to significantly improve next generation engineered T cell therapies and provides a framework for the design of entirely novel cellular therapeutics for oncology, autoimmune and regenerative medicine applications.

Past decades have witnessed the development of computational methods for creating novel protein domain structures and binding interactions^61–64^. However, protein allosteric functions such as signal transduction relying on long-range structural changes and dynamic communication between protein domains have largely been neglected and remain challenging to design. By leveraging fast protein structure and dynamics calculations, our computational platform provides a practical and efficient solution to the optimization of protein association and long-range mechanical coupling that govern signal transduction in single pass multi-domain membrane receptors. Based on general biophysical principles, the approach is not limited to a particular scaffold architecture or molecular mechanism and can be applied to design a wide range of biosensor functions. *De novo* globular domain structures and ligand binding domains can now be reliably designed using deep-learning based approaches^62,65–69^ and inserted as building blocks into our assembly protocol to couple any desired cues to arbitrary cell signaling. As such, our strategy should have a significant and far-reaching impact in basic and synthetic biology.

Splitting the T cell activation signals into separately expressed transgenic receptors allows for context-dependent tuning of T cell activation and anti-tumor activity and simultaneously enhances the tumor specificity of engineered T cells^70–72^. TME-derived IL4 or transforming growth factor β (TGF-β) have been used to elicit an immune stimulatory signal through empirically assembled chimeric receptors that leverage endo-domains from classical co-stimulatory (4-1BB) or cytokine receptors (IL7Rα)^73–76^. An engineered autocrine feedback loop based on GM-CSF, produced from T cells upon antigen recognition by the CAR, with a signaling output linked to IL18, has been explored as strategy to enhance anti-tumor responses of CAR T cells^77^. Constitutively active cytokine receptors have also been explored in combination with CARs, but those do not provide context-dependent functions^78,79^. Despite sustained efforts to develop chimeric receptors exploiting TME associated input signals to enhance CAR T cell therapy outcome, only a few chimera responding to soluble factors with demonstrated advantage in preclinical studies have been reported so far^20,73–76^. In contrast to these traditional empirical domain swapping chimeric designs, our approach highlights the benefits of in silico pre-screening, creating non-intuitive receptor scaffolds that would otherwise never be identified. Our approach also avoids extensive experimental screens, focusing experimental validation on selecting the best combinations of ligand binding, constitutive activity, and dynamic signaling responses. Given the numerous factors affecting multidomain signaling receptor structure and function, computational approaches like ours can significantly accelerate the engineering of receptors with customized functions, addressing a critical challenge in synthetic biology.

Based on rational design principles of signal transduction, our technology can engineer synthetic chimeric receptor structures with predictable and desired signaling output and sets the stage for the broader and more efficient development of biosensors with novel input-output functions. We validated our approach by building two distinct classes of chimeric signaling receptors that have only the TM and CT regions in common. In fact, the extracellular parts of VEGFR2 and CSF1R share very little sequence and structure homology and have no binding motifs in common. For each of these two classes, we have created several variants with rationally tuned constitutive and ligand-induced activities by programming the two most important features of receptor engineering, i.e. the combination of extracellular domains and the type of inter-domain linkers. Therefore, our study constitutes a strong proof of concept of the rational bottom-up assembly and design of chimeric receptors and supports the generalizability of the engineering approach. Since multiple methods can now confidently model a large variety of TM sequences and structures^65,80–85^, we think that our approach could also be applied to the design of chimera involving alternative TM and CT sequences. Lastly, our approach is not restricted to specific natural protein domains but could in principle assemble fully synthetic components provided their structure can be reliably predicted.

The potential field of application of T-SenSERs is large. Here we performed extensive experimental characterization when T-SenSERs are co-expressed with CARs in T cells. However, we envision future applications of VMR or CMR in other therapeutic T cell products, such as CAR-T cells with other endo-domains, TCR-T cells, tumor infiltrating lymphocytes or virus-specific T cells, where significant room for improvement exists, and TME specific enhancements of the anti-tumor response could be beneficial. Beyond αβ-T cells as cellular therapy platform, other cell types should also be investigated, for example γδ-T cells, NK cells or cytokine induced killer cells, that are amenable to off-the-shelf allogeneic cellular therapy development. The versatility of our computational platform will also allow the efficient adaptation of T-SenSER signaling domains to other cellular contexts, and to the exploitation of other disease specific inputs when envisioning applications in the autoimmune disease or regenerative medicine fields.

In conclusion, we have developed a novel in silico method for the assembly and design of synthetic receptors with programmable signaling activity for therapeutic applications. We demonstrate that these novel synthetic receptors can be tuned to enhance CAR-T cell functions by providing weak constitutive and/or strong ligand-specific co-stimulatory and cytokine signals upon sensing soluble factors enriched TMEs. We fully characterized two selected T-SenSERs in two completely independent experimental systems in combination with various CARs targeting both solid tumors and hematologic malignancies. In principle, T-SenSERs can be expressed in any engineered cell therapy for cancer or other diseases, in various immune cell types, activating pathways that favor long-term persistence and function of immune effector cells that directly eliminate or support the elimination of cancer cells. In addition to soluble factors, our technology can also be broadly applied to the sensing of chemical or mechanical cues such as cell surface ligands, extracellular matrix components or metabolites. Overall, the ability to engineer and control signal transduction should significantly impact basic and translational cell biology, synthetic biology and biomedicine.

## Data availability

The authors declare that all data supporting the findings in this study are either presented within the article or available from the corresponding author upon reasonable request.

## Acknowledgements

We are very grateful to Dr Stephen Gottschalk, Baylor College of Medicine and St. Jude Children’s Research Hospital, for providing the previously published EphA2 CAR constructs and the A549.GFP-FFluc cell line. We thank Rajendra Sharma (Barth lab) for initiating the computational framework, and Maude Varrin (Arber lab) for technical assistance and lab management (you left too early, RIP). We also thank Daniel Meraviglia, Lenka Polak, Patrick Reichenbach and Romain Vuillefroy de Silly for technical support, and Greta Giordano Attianese and all Arber and Barth lab members for helpful discussions and comments.

## Funding

J.A.R. was supported by a Swiss Government Excellence Scholarship for Foreign Scholars and the Emma Muschamp Foundation. P.B. is supported by a Swiss National Science Foundation grant (SNSF grant 31003A_182263), Swiss Cancer Research (KFS-4687-02-2019), funds from EPFL and the Ludwig Institute for Cancer Research. C.A. received funding for this study from Swiss Cancer Research KFS-4542-08-2018-R, Stiftung für Krebsbekämpfung, and the Lausanne University Hospital.

## Author contributions

J.A.R. and N.N. designed research, performed experiments, analyzed and interpreted results, and wrote parts of the manuscript. L.S.P.R. and A.F. designed research, developed the de novo assembly method for modeling and design of biosensors. L.S.P.R. analyzed and interpreted results, wrote and released the software, and wrote parts of the manuscript. A.F. designed research, developed the de novo assembly method for modeling and design of biosensors, analyzed and interpreted results. A.C. performed the mechanical coupling calculations and analyzed results. J.A.R., T.Q., C.V.G., C.P., F.B. and Y.B. performed experiments and analyzed results. P.B. conceived the computational study, designed research, supervised the study, analyzed and interpreted results and wrote the manuscript. C.A. conceived the study, designed research and supervised the entire study, performed experiments, analyzed and interpreted results and wrote the manuscript. All authors reviewed and approved the final version of the manuscript.

## Competing interests

C.A. and P.B. hold patents and provisional patent applications in the field of engineered T cell therapies. J.A.R. and L.S.P.R. hold a provisional patent application in the field of engineered T cell therapies. C.A. receives licensing fees and royalties from Immatics (through previous institution Baylor College of Medicine), participated in advisory boards for Kite/ Gilead, Janssen and Celgene/ BMS, received sponsored travel from Gilead (through current institution University Hospital Lausanne). All other authors declare no competing financial interest.

